# Sequence alignment using large protein structure alphabets improves sensitivity to remote homologs

**DOI:** 10.1101/2024.05.24.595840

**Authors:** Robert C. Edgar

## Abstract

Recent breakthroughs in protein fold prediction from amino acid sequences have unleashed a deluge of new structures, raising new opportunities for expanding insights into the universe of proteins and pursuing practical applications in bio-engineering and therapeutics while also presenting new challenges to protein search and analysis algorithms. Here, I describe Reseek, a protein alignment algorithm which improves sensitivity in protein homolog detection compared to state-of-the-art methods including DALI, TM-align and Foldseek, with improved speed over Foldseek, the fastest previous method. Reseek is based on alignment of sequences where each residue in the protein backbone is represented by a letter in a novel “mega-alphabet” of 85,899,345,920 (*∼* 10^11^) distinct states. Code is available at https://github.com/rcedgar/reseek.

## Introduction

Living cells are teeming hives of bustling molecular machines. The hive is constructed and directed by instructions encoded in nucleic acid chains: chromosomes and their delegates such as messenger RNAs. The machines themselves are built from proteins, with rare but notable exceptions including RNA subunits of the ribosome. Understanding, manipulating and designing proteins is a foundational project of molecular biology, where forensic traces of evolution provide vital clues for inferring function and mechanism. Vast collections of protein amino acid (a.a.) sequences have been generated by low-cost next-generation generation sequencing, leading to millions of high-quality structures enabled by recent breakthroughs in fold prediction. Exploiting this new trove of data raises challenges for computational methods developed in a previous era where solved structures numbered in the tens of thousands. Fundamental computational tasks include detection and alignment of homologous proteins, followed by identification of residue function via inferences of constraints on acceptable mutations made from such alignments.

### Pair-wise alignment and search

Quantitative evidence of relatedness can be obtained from a pair-wise protein alignment *a* by calculating a numerical value generically called an alignment score, here denoted *s*(*a*). Simple examples of scores include the fraction of identical a.a.s at corresponding positions (so-called identity) and the root mean square distance (RMSD) between corresponding *C_α_* atoms. Call a protein of interest the query (*Q*), a collection of structures the database (*D*), and a given protein in the database the target (*T*). A search is executed by constructing pair-wise alignments *{a*(*Q, T*) *∀ T ∈ D}*. Alignments are ranked by score to sort (hopefully) homologs above unrelated structures. Popular pair-wise protein structure alignment algorithms include DALI (Holm and Sander, 1993), TM-align (Zhang and Skolnick, 2005) and Foldseek (Van Kempen et al., 2024). DALI has the highest homolog recognition accuracy on recently-published benchmark tests (Holm, 2019; Van Kempen et al., 2024), while Foldseek has high reported accuracy at speeds thousands of times faster than TM-align and DALI, sufficient to enable a free public web service for searching large databases of predicted folds (https://search.foldseek.com).

### Alignment scores and E-values

For accurate homolog recognition, alignment scores of related pairs of proteins should tend to be higher than unrelated proteins. In practice, a biologist must choose where to draw the line between putative homologs and unrelated proteins by setting a cutoff, making an unavoidable trade-off between high sensitivity (maximizing the number of true homologs) and high specificity (minimizing the number of false positives). The gold standard approach to informing the choice of cutoff, pioneered by BLAST (Altschul et al., 1990), is to calculate a so-called E-value from the score *s*(*a_Q,T_*), i.e. an estimate of the expected number of false positives that would be obtained by setting the cutoff at this score. In other words, if an algorithm reports E-value *E_Q,T_* for its alignment of query *Q* to target *T ∈ D*, this can be regarded as the algorithm’s estimate of the mean number of false-positive errors per query (*FPEPQ*) that will occur if searches of *D* are executed for a representative selection of queries, and alignments with *E ≤ E_Q,T_* are considered to be predictions that the proteins are homologous. *FPEPQ* is sometimes called *EPQ* in the literature; my notation emphasizes that false-negative errors are not included. Biologists typically choose a cutoff *E_max_ <<* 1, implying a low false-positive probability. While the score depends only on the alignment, its E-value must additionally depend on the database *D*. In particular, if the number of database proteins *N* = *|D|* increases, the E-value should also increase. This follows because the expected number of false positives can be estimated as *E ≈ N × P*(*U|Q, s*), where *P*(*U|Q, s*) is the probability that a randomly-chosen unrelated protein aligns to *Q* with score *≥ s*, on the assumption that almost all database sequences are unrelated. Given a procedure for randomly sampling proteins, *P*(*U|Q, s*) is a function of *Q* and *s* independent of the database, and under these assumptions *E* is linearly proportional to *N*. DALI and TM-align report pair-wise scores, denoted Z and TM respectively, but not E-values. Rules of thumb have been suggested by the authors of these algorithms, e.g. *Z >* 4 or *TM >* 0.6 are good indications of homology, but these guidelines were developed when available structures numbered in thousands rather than millions, and it is unclear *a priori* how to adjust these rules when databases are large. Foldseek reports an E-value, and this E-value was shown to be effective in sorting homologs from non-homologs, but its accuracy as an estimate of *FPEPQ* was not investigated.

### Assessment of homolog detection accuracy

The SCOP database (Murzin et al., 1995) classifies proteins into folds, superfamilies and families. Superfamilies are believed to be homologous, while different superfamilies in the same fold are considered most likely to be similar due to convergence rather than homology. Families are somewhat arbitrary sub-divisions of superfamilies. The *de facto* standard benchmark for assessing protein homolog recognition accuracy is SCOP40, a subset of SCOP such that the maximum a.a. sequence identity is 40%. SCOP annotates domains, i.e. contiguous segments which are believed to fold independently and to be inherited intact from a single common ancestor. While roughly half of well-characterized proteins comprise a single domain by this definition, the remainder are concatenations of two or more (Ekman et al., 2005). For the work reported here I used SCOP v1.75, an older version of SCOP which uses the same domain identifier scheme as the Foldseek supplementary data to enable direct comparison. SCOP40 v1.75 has 11,206 domains classified into 1,194 folds, 1,960 superfamilies and 4,161 families. By contrast, the AlphaFold Protein Structure Database (APSD) (Varadi et al., 2022) contains *|APSD| ≈* 200M structures at the time of writing, i.e. *∼* 20, 000*×* more than SCOP40, suggesting that accuracy measured on SCOP40 may need substantial adjustments to extrapolate to database sizes used in practice. At a given cutoff, the numbers of true positives (*TP*), false positives (*FP*), true negatives (*TN*) and false negatives (*FN*) are calculated, and accuracy is summarized by the true positive rate *TPR* = *TP/NR* (also called sensitivity or coverage), false positive rate *FPR* = *FP/*(*TP* + *FP*), and *FPEPQ* = *FP/N*. Accuracy for a range of cutoffs can be summarized by a sensitivity vs. error plot (Brenner et al., 1998) with *TPR*=sensitivity on the *x*-axis and *FPEPQ* or *FPR* on the *y*-axis. For the results reported here, trivial self-alignments were excluded.

### Sensitivity up to the first false positive

The Foldseek paper (Van Kempen et al., 2024) uses sensitivity up to the first false positive (which I will call *SFFP*) as its preferred measure of sensitivity. Alignments for each query are considered separately, sorted by decreasing score or increasing E-value, and the number of hits above the first false positive is counted. Sensitivity is then calculated as the total number of retained alignments divided by the number of related pairs in the database, ignoring trivial self-alignments. In effect, this measures sensitivity assuming that the optimal cutoff for each query is known. The captures the effectiveness of an algorithm’s score or E-value in ranking related proteins above unrelated proteins, but surely overestimates the sensitivity that can be achieved in practice because a cutoff must be chosen, and the (unknown) optimal cutoff varies between queries.

### Modeling realistic searches

*FPR* should be approximately independent of the database size *N*, assuming that databases are sampled from the same underlying distribution, but *FPEPQ* is proportional to *N*. This raises the question of which regime of a sensitivity-error plot for SCOP40 is relevant to practical searches on databases with hundreds of millions of proteins. When using popular search tools such as BLAST or HMMer (Eddy, 2011), biologists typically choose an E-value threshold *E_max_ <<* 1 which implies a low estimated probability that the alignment is a false positive. E-value is an estimate of *FPEPQ*, suggesting that sensitivity should be measured with *FPEPQ <<* 1 to reflect the sensitivity that would typically be obtained in practice. Thus, to estimate sensitivity on APSD with a score cutoff giving *FPEPQ <* 1, sensitivity would ideally be measured at an *FPEPQ* value of *<* 1*/*20, 000 on SCOP40. However, obtaining *FPEPQ* values much less than one requires setting a cutoff such that false-positive errors occur in only a small subset of queries. With a small database such as SCOP40, the sub-sample will contain few queries and yield a noisy estimate. Here, I report sensitivity as the true-positive rate at *FPEPQ* values of 0.1, 1 and 10 (*Sens*(0.1), *Sens*(1) and *Sens*(10), respectively), noting that this approach may drastically overestimate sensitivity that could be achieved in practical searches of a much larger database such as APSD.

### Objectives for structure alignment

The conceptually simplest approach to aligning two protein structures is to seek a rigid-body transformation of one backbone which minimizes the RMSD of corresponding *C_α_* atoms (Kabsch, 1976; Umeyama, 1991). This works well for closely-related proteins. However, for more distantly-related proteins, three problems arise: (1) similarity may be limited to a sub-segment of one or both structures so that local rather than global alignment is required, (2) some regions may “flex” relative to others over evolutionary time so that somewhat different transformations should be applied to different segments, and (3) the number of *C_α_* atoms varies due to insertion and deletion mutations causing pair-wise correspondences to become unclear (for example, choosing the closest *C_α_* may result in asymmetrical or many-one correspondences). Many different objective functions have been proposed for optimizing local alignments of *C_α_* coordinates, including the DALI Z-score, TM-score and CE (Shindyalov and Bourne, 1998). An alternative approach is to represent a structure as a sequence of letters, one for each *C_α_*, and adapt well-established local sequence alignment algorithms such as Smith-Waterman (Smith et al., 1981) and BLAST. This approach was pioneered by 3D-BLAST (Yang and Tung, 2006) and CLePAPS (Wang and Zheng, 2008), which used conformational alphabets with 23 and 17 letters, respectively. In this type of alphabet, a letter represents the local conformation (secondary structure) of a *C_α_* ; for example, if the *C_α_* is in an alpha helix its letter might be ‘H’. Local conformation can be described by real-valued features including angles and distances which are invariant under rigid translation and rotation, i.e. independent of the arbitrary choice of axes for *C_α_* coordinates (Hovmöller et al., 2002). Features for a reference set of *C_α_* s are clustered to obtain a representative subset, and a letter for a given *C_α_* is assigned by identifying the closest representative. In this way, continuous parameters characterizing local conformation are condensed into a discrete alphabet *A* with *|A| ∼* 20 letters. The score of an alignment is the sum of log-odds scores for aligned pairs of letters, minus gap penalties, where the symmetrical *|A|×|A|* matrix *M^A^* of scores for every possible pair of letters is pre-trained on a reference set of trusted alignments. Training of this matrix uses methods developed for a.a. substitution matrices such as BLOSUM62 (Henikoff and Henikoff, 1992). Foldseek extends this approach from secondary to tertiary structure by considering the conformation of a *C_α_* and its nearest neighbor *C_α_* by Euclidean distance, excluding adjacent positions in the backbone. Their combined conformation is represented by a 10-dimensional feature vector and condensed into a 20-letter alphabet called 3Di. The score of an aligned pair of *C_α_* s is calculated as a weighted sum of the log-odds score for their 3Di letters and the BLOSUM62 log-odds score of their corresponding a.a.s. The relative weight of scores for 3Di and a.a. letters is a constant parameter of the algorithm fixed by training on a reference set.

### Structure alphabet size

Intuitively, it seems obvious that a great deal of information must be lost when backbone coordinates are condensed into a sequence of letters from an alphabet of size *∼* 20 letters. Consider Foldseek’s 3Di, which represents the conformation of a *C_α_* and its nearest neighbor. Secondary structure can be condensed to an alphabet with four states *{*helix, strand, turn, loop*}*, which already loses information about beta strands (e.g. are they parallel or anti-parallel?), and lumps a rich variety of loop conformations into a single state. With one residue plus its neighbor, this gives 4 *×* 4 = 16 states, leaving little room to capture much more information in a 20-state alphabet. The Foldseek authors investigated the relationship between alphabet size and sensitivity in their framework. Their supplementary Fig. 13 shows that a point of diminishing returns was reached at *∼* 20 letters. I believe the diminishing returns reflects increased difficulty in estimating the scoring matrix from a relatively small training set rather than an inherent limit to the power of larger alphabets. The log-odds score 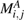 for aligning letter *i* to letter *j* in alphabet *A* is calculated as

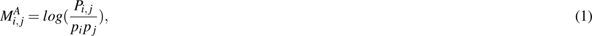

where *P_i, j_* is the estimated joint probability in the training set of (*i, j*) as a fraction of all aligned pairs, and *p_i_* is the probability of letter *i*. Probabilities are estimated as the observed frequencies. The probability distribution *p_i_*is likely to be skewed with a tail of rare letters. With larger alphabets, the numbers of observed aligned pairs of rare combinations become too small to obtain an accurate estimate of joint probabilities *P_i, j_, i ̸*= *j* for low-frequency letters. Reseek solves this problem, thereby enabling arbitrarily large structure alphabets to be deployed in practical applications, by factoring structure states into tractable components whose log-odds scores are calculated independently.

### Reseek structure representation

Reseek represents a *C_α_* backbone atom and its structural context as a feature vector (FV) which may have many components. A component may be single real value (scalar), multiple real values (vector), or inherently discrete. Often, one component is the amino acid type, an inherently discrete feature which can be represented by the standard 20-letter alphabet. Examples of real-valued features include *DistNEN*, the distance in Angstroms between the *C_α_* at position *i* and its nearest Euclidean neighbor (*NEN_i_*, i.e., the neighbor as defined by Foldseek), and *DistREN*, defined as the distance to the nearest neighbor in the opposite direction in the chain compared to its NEN (its reverse Euclidean neighbor, *REN_i_*) as shown in Fig. 1. The local conformation (*Con f*) at position *i* in the backbone is represented by real-valued features comprising all-vs-all pair-wise distances between the 2*κ* + 1 positions centered at *i*, excluding adjacent pairs which are at an effectively constant distance set by the *C_α_ −C_α_* bond length. For the results reported here, *κ* = 3. Similarly, the local context around the NEN and REN for position *i* is represented by all-vs-all pair-wise distances between the 2*κ* + 1 positions centered at *NEC_i_* and *REC_i_*, respectively. For alignment construction and scoring, the FV is condensed into a discrete feature vector (DFV) in which each component is a letter from an alphabet which is typically chosen to have size ≲ 20 to mitigate problems with sparse training data for low-frequency letters. The procedure for condensing a feature depends on whether it is a scalar or vector.

**Figure 1.**
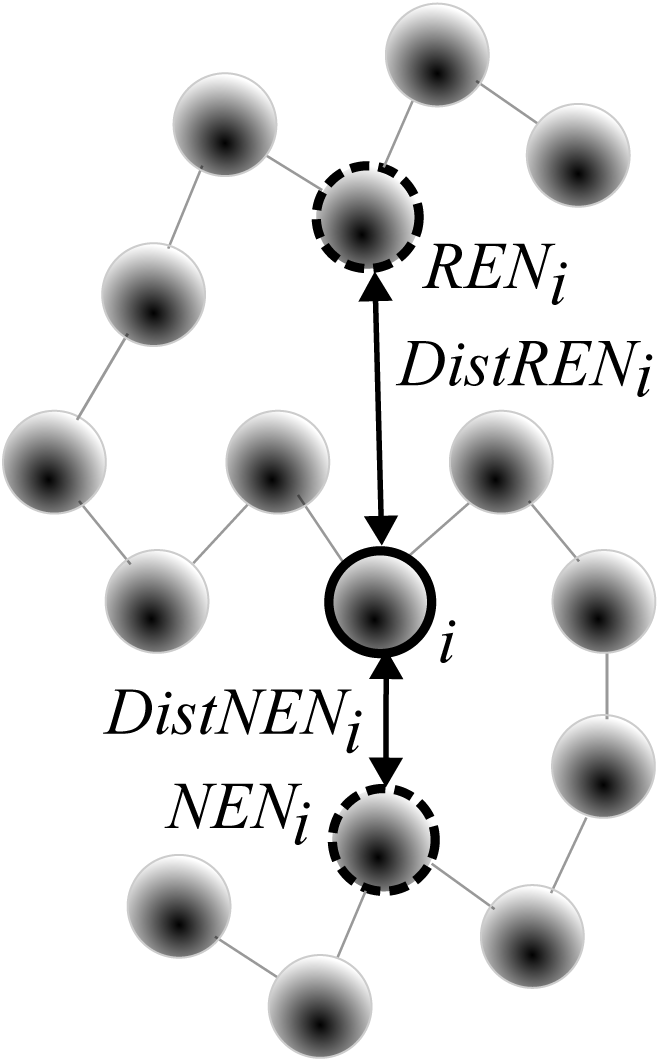
Nearest-neighbor distance features. Shaded spheres represent *C_α_* atoms, lines represent *C_α_ −C_α_* bonds forming the protein backbone. Position *i* is indicated by a solid circle, its nearest neighbors by dashed circles. The nearest Euclidean neighbor (*NEN_i_*) is the closest *C_α_* atom to *i*, ignoring adjacent residues in the chain. The reverse Euclidean neighbor (*REN_i_*) is the nearest neighbor in the reverse chain direction compared to *NEN_i_*. Feature *DistNEN_i_* is the distance in Angstroms between *i* and *NEN_i_*, similarly *DistREN_i_*is the distance to *REN_i_*.

### Condensing a scalar feature

For the prototype implementation of Reseek which generated the results reported here, scalar features were condensed as follows. Feature values for all *C_α_* s in the training set are collected and sorted by increasing numerical value. If the desired alphabet size is *L* letters, the sorted list is divided into *L* contiguous bins of equal size, and *L−* 1 threshold values are calculated which divide the bins. To achieve this, the threshold *t_B_*between bin *B* and bin *B* + 1 is set to the mean of the maximum value in bin *B* and the minimum value in bin *B* + 1. The condensed alphabet letter of feature value *x* is then *argmin*(*B* : *x > t_B_*), i.e. the lowest bin for which *x* is above the threshold. Thus, letter assignment is reduced to comparison with *L* thresholds. This procedure is designed to ensure that letters have approximately equal frequencies, maximizing the entropy of the alphabet and thereby maximizing the information in a sequence (Strait and Dewey, 1996).

### Condensing a vector feature

For the prototype implementation of Reseek which generated the results reported here, vector features were condensed as follows. Vectors were collected from all *C_α_* s in a training set of structures and clustered by *K*-means (MacQueen et al., 1967), where *K* was set to the desired alphabet size *L*. Following training and storage of the cluster representatives, the letter for a feature vector is assigned by finding the closest representative. For search, this procedure for assigning letters may be more efficient than the corresponding procedure for the 3Di alphabet, which requires a neural network classifier at runtime.

### Training the log-odds score matrix for discrete features

Log-odds score matrices for discrete features, including the amino acid type, are calculated per Eq. 1, where probabilities are estimated as the frequencies observed in a training set of alignments. For the work reported here, the training set was constructed using TM-align from a subset of SCOP40 pair-wise alignments with 0.6 *< TM <* 0.8. This range was chosen by intuition driven by experience with BLOSUM matrices, where BLOSUM62 has been found to be the best matrix for general-purpose homolog search using a.a. sequence (Altschul et al., 1997; Styczynski et al., 2008). BLOSUM62 was trained on alignments with *>* 62% identity, and intuitively 0.6 *< TM <* 0.8 seemed to be a comparable choice for structural alignments, i.e. intermediate between high-similarity alignments where any reasonable matrix works well and a challenging regime where similarity is low and training alignments may be dubious.

### Log-odds score for a pair of DFVs

As described above, the state of a *C_α_* atom is condensed into a discrete feature vector (DFV). Let the vector for the *C_α_* at position *i* in a given protein be *V* (*i*), with integer-valued components (letters) *V_f_* (*i*)*, f ∈* **F** where **F** is the chosen set of features. The score for an aligned pair of positions *i, j* is calculated as

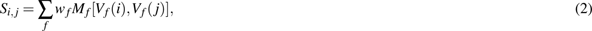

where *M_f_* is the log-odds score matrix for feature *f*, and *w_f_* is its relative weight. A DFV has

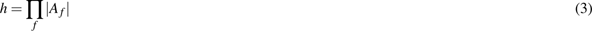

possible states, where *A_f_* is the alphabet for feature *f*, and a given DFV may therefore be regarded as a letter in a “mega-alphabet” **H** with *h* “mega-letters”. From this perspective. Eq. 2 is an estimate of the log-odds score for a pair of mega-letters that would be obtained with a sufficiently large training set.

### Local alignment construction

The score of an alignment of proteins *Q* and *T* is

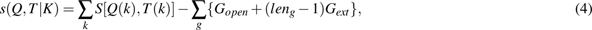

where *K* is the set of columns in the alignment, *k ∈ K* is a column, *Q*(*k*) is the position in *Q* that appears in that column, *T* (*k*) is the position in *T*, *S* is calculated according to Eq. 2, *g* is a gap in the alignment, *len_g_* is the number of columns in the gap, *G_open_* is the gap-open penalty and *G_ext_* is the gap-extension penalty. A high-scoring local alignment is found using an algorithm such as Smith-Waterman or a BLAST-like method which seeks to optimize *s* as given by Eq. 4.

### Contact map similarity

The contact map of a protein is a symmetrical matrix giving all-by-all pair-wise Euclidean distances between its *C_α_* s. This matrix is often visualized as a “heat” map (closer is warmer), which enables visual identification of domain architecture, secondary structure and contact clusters (Vehlow et al., 2011). Structures can be aligned by aligning their contact maps, as pioneered by DALI, which attempts to maximize the following heuristic score:

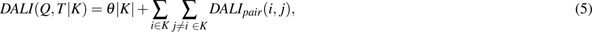

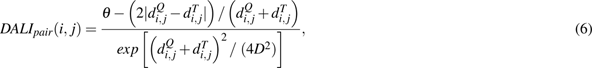

where *i* and *j* are columns in the alignment, *|K|* is the number of columns, *θ* = 0.2 Angstroms (Å), *D* = 20 Å, and 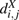 is the Euclidean distance in Å between the *C_α_* s from protein *X* found in columns *i* and *j* respectively. Given an alignment, the DALI score can be calculated in *O*(*N*^2^) time and space for proteins of length *N*, while constructing an alignment by exact optimization of the DALI score is NP-hard (Wohlers et al., 2012).

### Reseek test statistic and E-value

A test statistic is a numerical value calculated from an observation which is compared to a null model distribution to assess the significance of the observation, i.e. to ask whether it is likely that this value could be obtained by random sampling from the chosen null model. For protein sequence alignment, the canonical example of a test statistic is the bit score of a BLAST alignment (Altschul et al., 1990), where the null model is alignment of the query to a random sequence. In the prototype described here, Reseek uses the following test statistic:

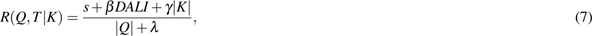

where *s* is given by Eq. 4, *DALI* by Eq. 5, *β* weights the contribution of *DALI*, *|Q|* is the length of the query protein *Q*, *λ* is a parameter introduced to damp the growth in *R* for anomalously short queries, and *γ* is a parameter introduced to reward longer alignments. The term *θ|K|* in *DALI* already rewards longer alignments, but empirically setting *γ* = 0.1 was found to improve accuracy. For the results reported here, *λ* = 32. The null model is defined as choosing an unrelated protein at random. An E-value is obtained from *R* by fitting to an empirical null-model distribution.

### Features

The Reseek framework accommodates an arbitrary number of structure features. This section summarizes features implemented in the prototype; see also Reseek structure representation above. Here, *i* is the position of the *C_α_* in the chain. If a feature is condensed to an alphabet with size *L*, it is denoted as *FeatureName*(*L*).

*AA* is the amino acid type, taking 20 possible values.

*DistNEN* is the distance in Angstroms between the *C_α_* and its nearest Euclidean neighbor (NEN). “Euclidean” means 3D space, to emphasize the distinction from distance measured as number of residues along the chain (Fig. 1).

*DistREN* is the distance in Angstroms between the *C_α_* and its reverse Euclidean neighbor (REN), i.e. its nearest Euclidean neighbor considering only the opposite chain direction from its NEN (Fig. 1).

*Con f* (local conformation) is a vector feature with all pair-wise distances between *C_α_* s in the range (*i−κ*) *…* (*i*+*κ*), i.e. the contact map for 2*κ* +1 residues centered at *i*, with *κ* = 3 by default. This feature captures secondary structure; in particular, *Con f* (3) correlates well with *{ helix, strand, other }* annotations from standard methods (see (Andersen and Rost, 2003)), and is used to identify helix and strand elements for other features.

*NENCon f* is the *Con f* of the NEN.

*RENCon f* is the *Con f* of the REN.

*NormDens* is normalized density, a measure of how many residues are found in the Euclidean neighborhood surrounding the *C_α_*, with more distant residues down-weighted. Density (*Dens*) is calculated as follows:

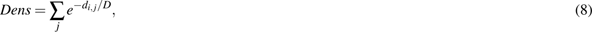

where *D* is the DALI *D* parameter (i.e., 20 Å), *d_i, j_*is the Euclidean distance between position *i* and position *j*, and *j* is summed over positions from (*i−* 50)*…*(*i* + 50) excluding positions (*i−* 3)*…*(*i* + 3) as close neighbors in chain order are certainly also close in Euclidean distance and therefore least informative. Normalized density (*NormDens*) is calculated by re-scaling *Dens* to a range zero to one, excluding outliers.

*HelixDens* is calculated by Eq. 8, including only positions which are alpha helices according to *Con f* (3).

*StrandDens* is calculated by Eq. 8,, including only positions which are beta strands according to *Con f* (3).

*DistNextHelix* is the Euclidean distance between position *i* and the mid-point of the nearest alpha helix later in the chain than position *i*, excluding the helix which contains *i* if any. Secondary structure elements are identified as contiguous segments with the same *Con f* (3). If there is no later helix, the distance is zero.

*DistPrevHelix* is similarly the Euclidean distance to the mid-point of the preceding alpha helix in the chain.

### Parameter training

The Reseek algorithm has a large number of parameters, including *G_open_*, *G_ext_*, *β*, *λ*, the features to include (**F**) as a subset of all implemented features (**F***^∗^*), and weights of the included features *{w_f_, f ∈* **F***}*. In the prototype described here, 11 features were implemented, giving 2*^|^***^F^***^∗|^* = 2^11^ = 2, 048 possible choices for **F**. For each possible **F**, there are *|***F***|* weights and four scalar parameters *G_open_*, *G_ext_*, *β* and *λ*. The following protocol was implemented to enable practical tuning of this large parameter space. A given subset of features and their proposed weights were first evaluated using training alignments by constructing a log-odds score matrix according to Eq. 1. Higher expected score of the matrix (*ES*, also known as relative entropy (Altschul, 1991)) was used as the objective. For a given subset and its weights, *ES* can be calculated from a large number of preconstructed alignments in a fraction of a second, enabling exploration of many parameter combinations. First, all pairs of features from **F***^∗^* were evaluated with a range of relative weights, keeping the best few feature pairs and their best weights, i.e. those with highest *ES*. For each such pair and its weights, all possibilities for adding a third feature were explored, again with a range of weights for the new feature. This procedure was iterated until a point of diminishing returns was reached by adding new features. This gave a pool of promising combinations for further consideration. The next stage was to evaluate candidate parameter sets by constructing ungapped alignments on a SCOP40 subset with high *SFFP* as the objective. Small variations in the weight of each parameter were tried one at a time, keeping changes that improved *SFFP*, repeating with increasingly small variations until convergence was reached. Finally, gapped alignments were generated to optimize *G_open_*, *G_ext_*, *β* and *λ* using a similar procedure to weights, i.e. by introducing variations one parameter at a time with high *SFFP* as the objective. Default parameters are shown in Table 1. The size of the default mega-alphabet is *h* = 20 *×* 16^8^ = 85, 899, 345, 920 letters.

**Table 1.**
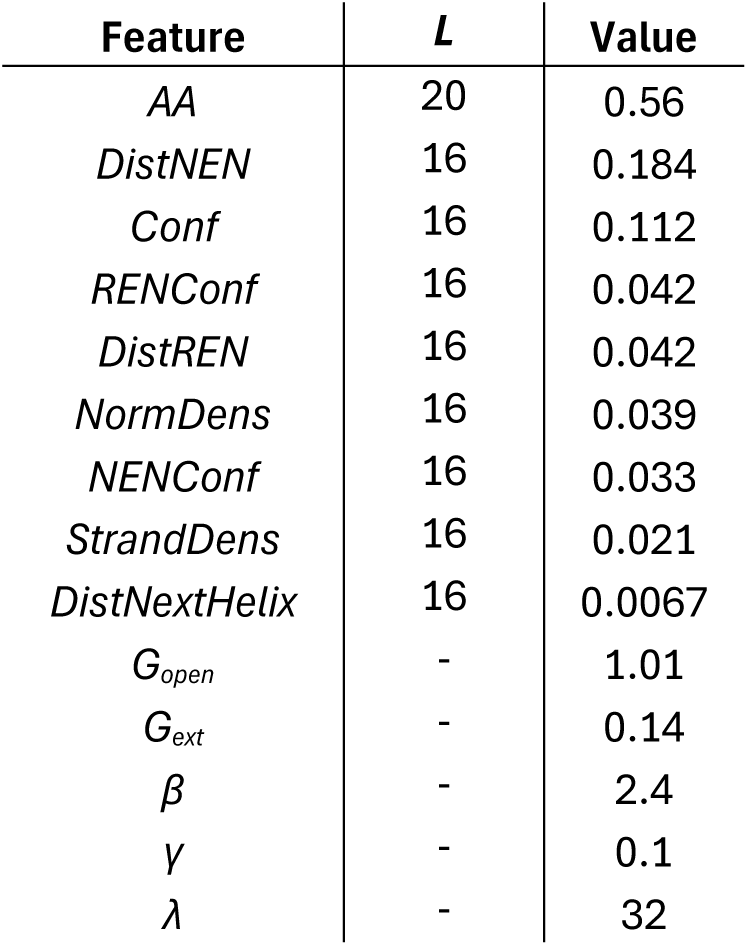
Default Reseek parameters. For a feature, *L* is the alphabet size and *Value* is its weight.

| Feature | $L$ | Value |
| --- | --- | --- |
| $AA$ | 20 | 0.56 |
| $DistNEN$ | 16 | 0.184 |
| $Conf$ | 16 | 0.112 |
| $RENConf$ | 16 | 0.042 |
| $DistREN$ | 16 | 0.042 |
| $NormDens$ | 16 | 0.039 |
| $NENConf$ | 16 | 0.033 |
| $StrandDens$ | 16 | 0.021 |
| $DistNextHelix$ | 16 | 0.0067 |
| $G_{open}$ | - | 1.01 |
| $G_{ext}$ | - | 0.14 |
| $\beta$ | - | 2.4 |
| $\gamma$ | - | 0.1 |
| $\lambda$ | - | 32 |

### MSP accelerator

The throughput of pair-wise comparisons is optionally accelerated by applying a filter before proceeding to construct the alignment required to calculate *R* per Eq. 7. The accelerator computes a similarity measure *ω* which should tend to be larger for more closely related structures:

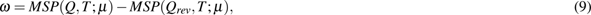

where *MSP* is the maximum segment pair score, *Q_rev_*is the chain of *Q* in reverse order, and *µ* is an alphabet chosen to balance the competing requirements of fast execution, which favors small alphabets, and effective filtering, which favors larger alphabets. *MSP* is the highest score which can be obtained by aligning a pair of contiguous sub-chains without gaps; it is calculated using a Smith-Waterman implementation with optimizations to exploit a scenario where the alphabet is small, both *Q* and *Q_rev_* are likely to be compared with many different targets, gaps are not allowed, and explicit alignments are not required, only scores. In the prototype, *µ* is the 36-letter alphabet *{Con f* (3)*, NENCon f* (3)*, DistREN*(4)*}*. A pair is discarded iff *ω <* Ω, where Ω can be chosen by the user. Smaller values increase speed, at the possible expense of reduced sensitivity.

### *K*-mer accelerators

Other accelerators are based on *k*-mers, i.e. sub-chains of fixed length *k C_α_* s, where a subset of *b ≤ k* letters may be considered according to a fixed pattern (Ma et al., 2002). *K*-mer indexes are widely used to accelerate searches in biological sequence applications (Marchet et al., 2021). The design of a *k*-mer index implies a complex trade-off between speed, sensitivity, and index size. To maximize speed, only exact *k*-mer matches (seeds) should be considered, because this enables the fastest lookups. The number of relevant exact seed matches may be optimized by varying *k*, *b*, the pattern of *b* letters within a seed, and the alphabet, and then seeking a design which maximizes sensitivity to related proteins while minimizing false positives. For U-sorting (Edgar, 2010), the correlation between E-value and number of unique seeds in common between pair of proteins should be maximized. Preliminary attempts were made to perform such optimizations for several types of seed index, all of which converged on the same seed design: *k* = 11, *b* = 2, *pattern* = 10000000001, with alphabet *µ* = *{Con f* (3)*, NENCon f* (3)*, DistREN*(4)*}* (i.e, the same alphabet as the MSP accelerator). This design has *|µ|^b^* = 1, 296 distinct seed values.

## Results

### Tested algorithms

I tested the following software. Reseek release v1.3-beta. DALILite v5 (August 2023). Foldseek release 8-ef4e960, which was the latest at the time these experiments were performed (April 2024). GTalign v0.15.0-alpha. TM-align was compiled from the latest available source at https://zhanggroup.org/TM-align/, which was dated 2022/4/12.

### SCOP40 tests of accuracy and speed

Sensitivity vs. FPEPQ on the SCOP40 test is shown in Table 2 and Figs. 2 and 3. *FPEPQ* = 10 corresponds to an ideal E-value threshold of 10 and therefore represents the high end of E-value thresholds typically used in practice, while sensitivities measured with smaller *FPEPQ* become increasingly noisy due to small numbers of errors. Superfamily (Fig. 2) is the preferred proxy for homology, where with *FPEPQ <* 10, Reseek has higher homolog sensitivity vs the state of the art represented by DALI, TM-align, GTalign and Foldseek.

**Figure 2.**
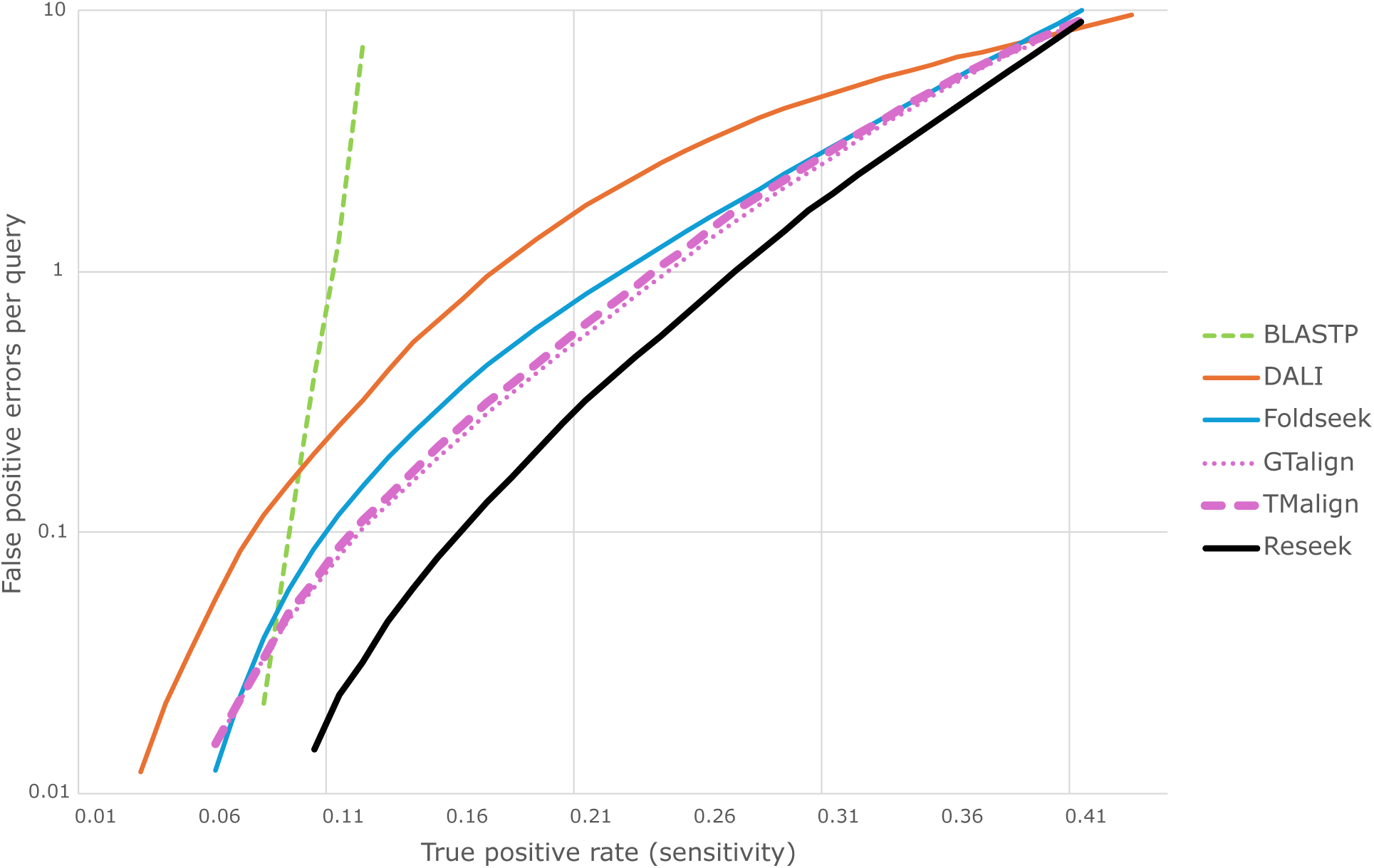
Sensitivity vs. errors for homology detection on the SCOP40 test. The *x* axis is true-positive rate (TPR, the fraction of all possible TPs). The *y* axis is false-positive errors per query (FPEPQ); note the log scale. Hits to the same (different) superfamily are considered true (false) positives. With decreasing *FPEPQ <* 1, measured FPEPQ becomes increasingly noisy due to the small sample size for errors. *FPEPQ* = 1 corresponds to an E-value threshold of 1 if E-values are correctly predicted. With *FPEPQ <* 10, Reseek achieves higher sensitivity compared to the state of the art represented by DALI, TM-align and Foldseek.

**Figure 3.**
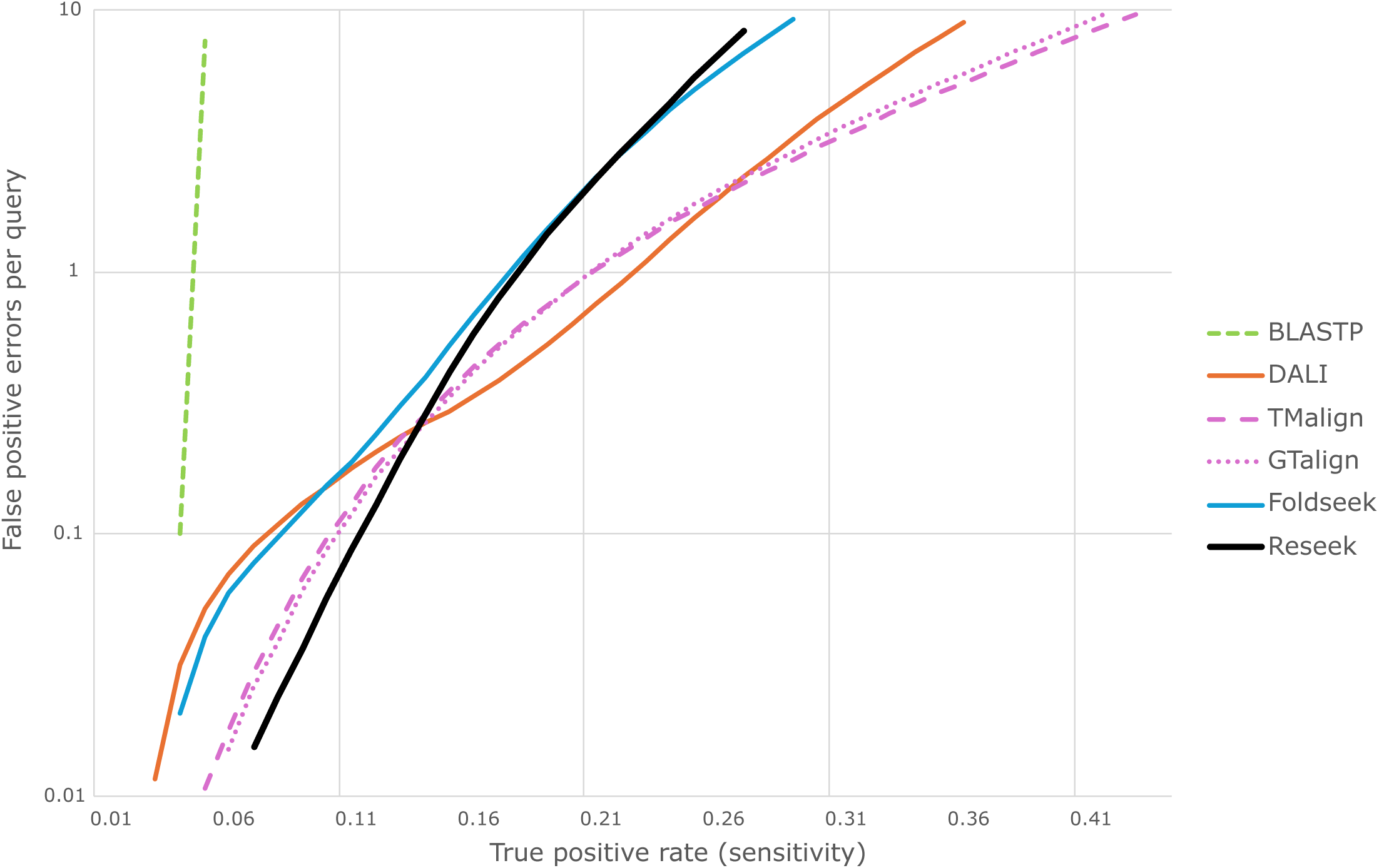
Sensitivity vs. errors according to SCOP fold. Analysis performed similarly to Fig. 2 except that fold instead of superfamily is the standard of correctness. Hits to different superfamilies in the same fold are most likely convergent rather than homologous. Here, Reseek has higher sensitivity below 0.1 errors per query but lower sensitivity above 1 error per query. I interpret the cross-over as reflecting an increasing number of convergent folds which are not homologous (see main text), noting that Reseek is explicitly designed to find homologs rather than convergent folds. The excess hits found by DALI, TM-align and GTalign with *FPEPQ >* 1 are then largely accounted for as having similar but non-homologous folds, which are likely to have radically different functions.

**Table 2.** Performance on the SCOP40 test. Time is wall-clock elapsed search time in seconds on a 3.2GHz Intel i9-14900K CPU (32 cores, running 32 threads or 32 parallel processes), excluding time to build the search database. *Sens*(*e*) is sensitivity at *FPEPQ* = *e*, *SFFP* is sensitivity up to the first false positive. *FF* is the fraction of non-singleton queries with a TP hit above the default threshold, *FF*1 is the fraction of *FF* hits that were above the first false positive, i.e. the fraction for which the top hit is in the same family. Foldseek is release 8-ef4e960, Foldseek(p) is data from the paper.

| Method | Time (s.) | Memory | Sens(0.1) | Sens(1) | Sens(10) | SFFP |
| --- | --- | --- | --- | --- | --- | --- |
| BLASTP | 127 | 35 Mb | 0.090 | 0.11 | 0.13 | 0.10 |
| DALI | 410,000 | 4.0 Gb | 0.075 | 0.17 | 0.44 | 0.49 |
| Foldseek | 78 | 1.8 Gb | 0.11 | 0.22 | 0.41 | 0.39 |
| GTalign | 173,000 | 6.0 Gb | 0.12 | 0.24 | 0.42 | 0.44 |
| TM-align | 370,000 | 175 Mb | 0.12 | 0.24 | 0.42 | 0.45 |
| Reseek | 68 | 319 Mb | 0.16 | 0.27 | 0.42 | 0.39 |

### Two-fold cross-validation

Reseek’s large parameter space raises potential concerns of over-fitting. To investigate this, I performed two-fold cross-validation by splitting families in SCOP40 into two equal-size subsets A and B. The parameters in Table 1 were trained separately on A and B, then used to measure *SFFP* and *Sens*(1) on both subsets. As seen in Table 3, results from self-training and cross-training are almost identical, showing that over-fitting is minimal. This justifies the use of default parameters trained on all SCOP40, which on the basis of these cross-validation results I would expect to generalize slightly better than parameters trained on a subset.

**Table 3.** Two-fold cross-validation of Reseek training on SCOP40. Families in SCOP40 were divided into two equal-sized subsets A and B. Parameters shown in Table 1 were optimized separately for A and B. This table reports *SFFP* and *Sens*(1) for all four possible combinations of training set and test set, showing that self-training gives almost identical results to cross-training.

| Test | Train | SFFP | SEPQ1 |
| --- | --- | --- | --- |
| A | A | 0.672 | 0.600 |
| A | B | 0.670 | 0.600 |
| B | B | 0.607 | 0.538 |
| B | A | 0.605 | 0.537 |

### E-value accuracy

E-value is a prediction of FPEPQ. Figs. 4 and 5 show E-value vs. FPEPQ for Foldseek release 8-ef4e960 and Reseek, respectively. The other tested methods do not report E-values. The Foldseek E-values are shown to substantially under-estimate error rates, while Reseek’s are in good agreement.

**Figure 4.**
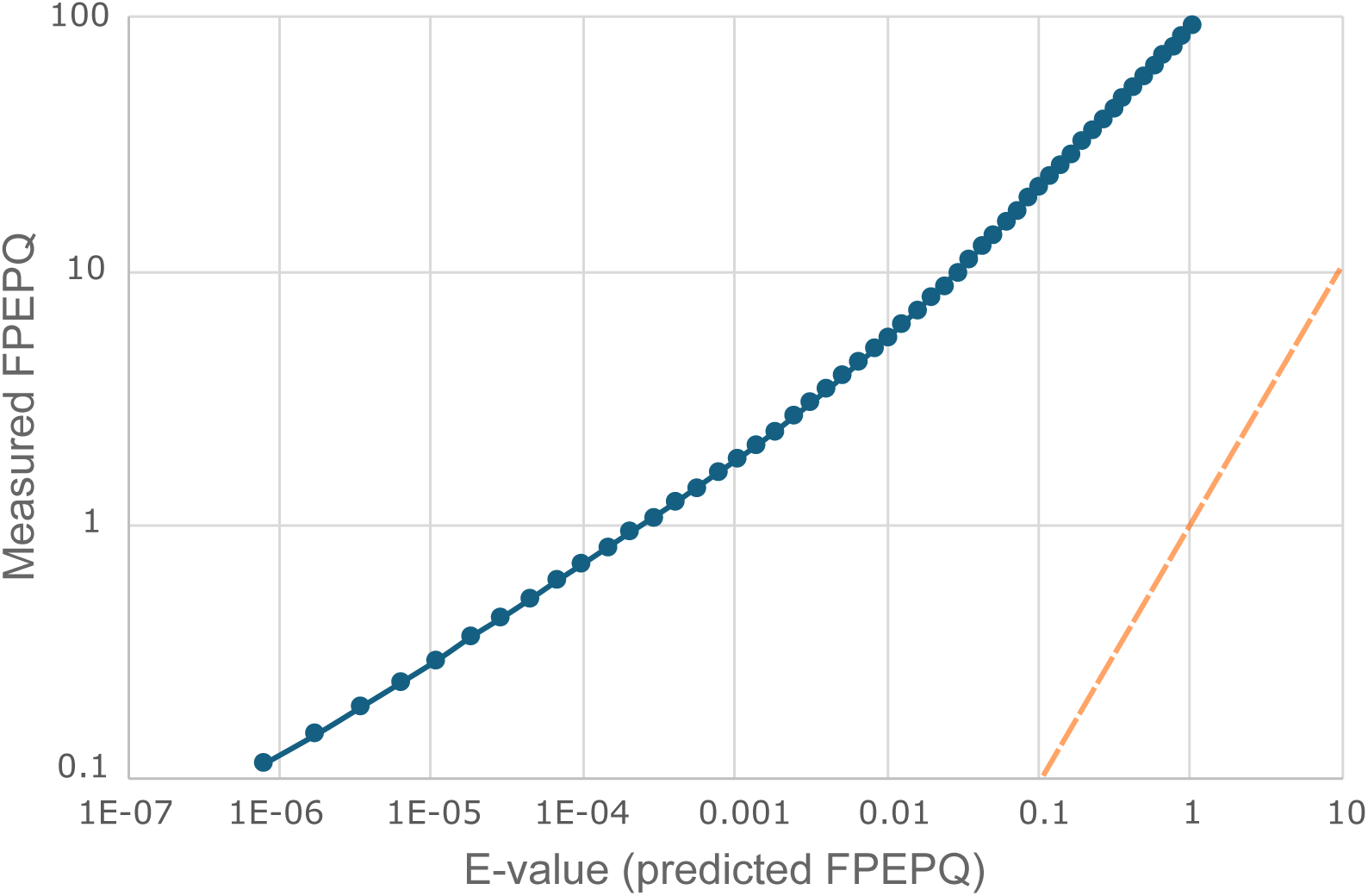
E-value vs. FPEPQ for Foldseek. E-value is a prediction of FPEPQ. This plot shows log-scaled E-value vs. FPEPQ on SCOP40 for Foldseek release 8-ef4e960 in the range *E* = 10*^−^*^7^ to *E* = 10, representing a range of thresholds that might typically be used in practice. The orange dashed line corresponds to a correct prediction of superfamily FPEPQ by the E-values, which is seen to radically depart from the correct values. For example, at an E-value threshold of 10*^−^*^4^, the measured FPEPQ is *∼* 1, i.e. *∼* 1, 000*×* higher than predicted. At E-value threshold 1, the measured error rate is *∼* 100. Thus, the Foldseek E-value greatly under-estimates the error rate on this test.

**Figure 5.**
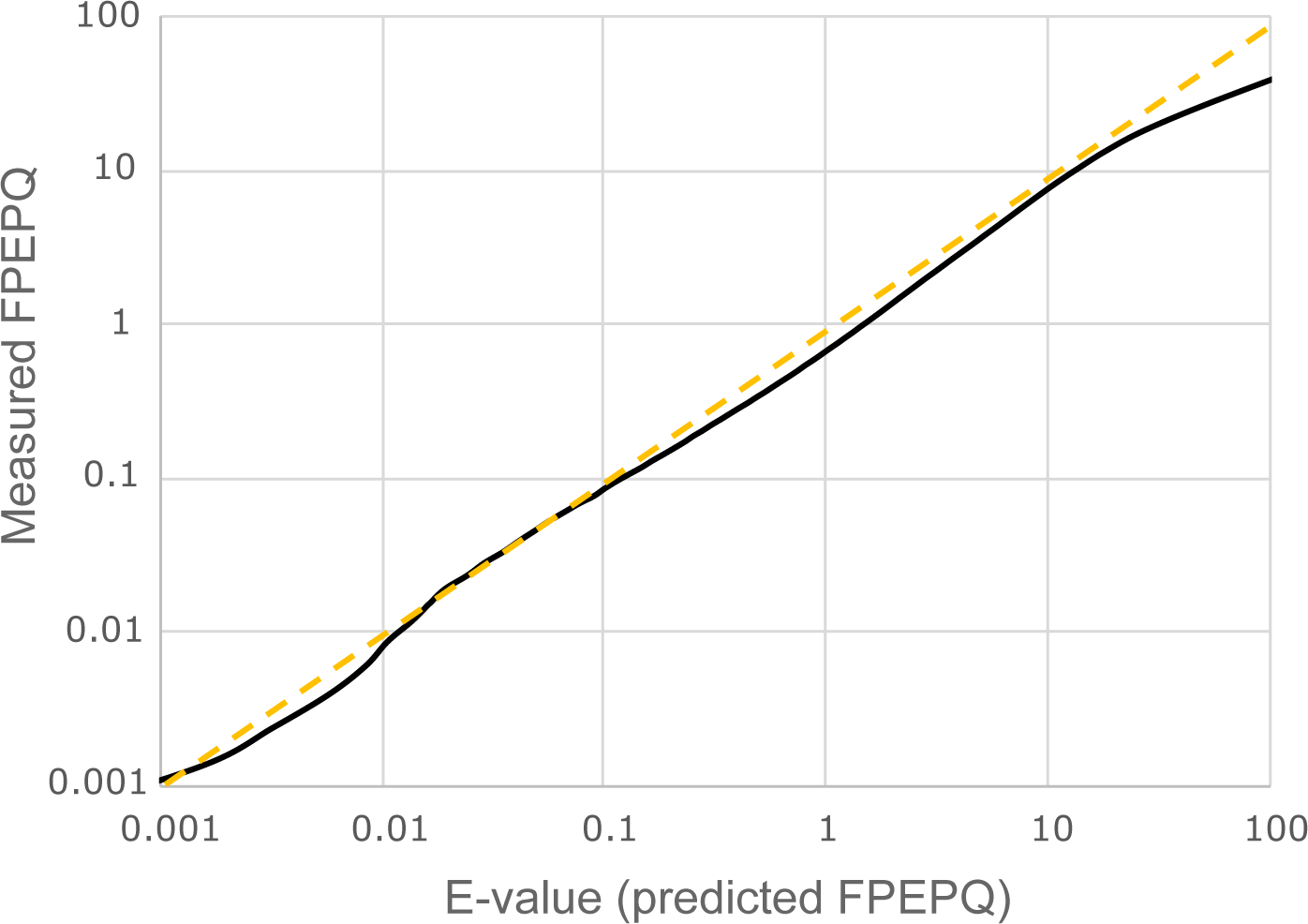
E-value vs. FPEPQ for Reseek. E-value is a prediction of FPEPQ. This plot shows log-scaled E-value vs. FPEPQ on SCOP40 for Reseek similarly to Fig. 4 for Foldseek. The orange dashed line corresponds to a correct prediction of FPEPQ by the E-values.

